# Synthesis and In Vitro Assessment of Triazole-Based Compounds as Potential Inhibitors of Herpes Simplex Virus Type 1

**DOI:** 10.1101/2025.05.03.652007

**Authors:** Neha U. Mishra, Kiran K. Sanap, Pritesh C. Sharma, Tulsiram Kudale, Jyoti Vanawe, Abhijit Biswas, Dhiraj Bhatia

**Author notes:** Correspondence to: Kiran K. Sanap-Shri Guru Gobind Singhji Institute of Engineering and Technology, Nanded, 431606, India, Abhijit Biswas − Department of Biological Sciences and Engineering, Indian Institute of Technology Gandhinagar, Palaj 382355, Gujarat India, Dhiraj Bhatia − Department of Biological Sciences and Engineering, Indian Institute of Technology Gandhinagar, Palaj 382355, Gujarat India.

## Abstract

A library of novel 1,2,4-triazole derivatives fused with a pyrazine moiety (5a–5t) was successfully synthesized and evaluated for their therapeutic potential against HSV-I virus and oxidative stress. These compounds were assessed for anti-inflammatory, antioxidant, and anti-HSV activities, demonstrating encouraging biological profiles. In particular, compounds 5f and 5t exhibited markedly improved antiviral efficacy compared to the reference drug, Acyclovir. The antioxidant capacity was also notable, with compound 5b showing exceptional radical scavenging potential. To better understand the interaction at the molecular level, docking studies were conducted, indicating that these molecules could potentially inhibit HSV1, a key enzyme required for viral replication. Also, in silico testing was conducted for assessing drug-likeness, ADME characteristics, and stability of metabolism which was helpful for optimizing further lead designs. Although the experimental methods were accurately implemented, some human error such as timing during administration of the compounds, measuring, and other tasks may have slight variations which are not entirely avoidable. These possible blunders are, however, unlikely to change significantly the observed trends. Ultimately, the study emphasizes these new hybrids of triazole-pyrazine as potential precursors for the synthesis of powerful anti-HSV-I and possibly antibacterial medicines, which require more advanced pharmacological and clinical research.

## 1. Introduction

Herpes simplex virus type 1 (HSV-1) remains one of the most common and enduring viral infections in humans It has been associated with various clinical disorders such as keratitis, oral and vaginal herpes, and in more severe forms, encephalitis.^1–3^ Despite the prevalence and routine use of antivirals, HSV-1 remains a significant public health concern due to its frequent occurrence and occasional reactivation. The infection’s recurrent nature predisposes the affected individuals to a host of capricious and disturbing symptoms, which result in a diminished quality of life. To manage these challenges, there is growing interest in focusing on alternative approaches that go beyond inhibiting viral replication, including targeting inflammation and oxidative stress. Researchers have recently turned to the development of new compounds with multi-targeted mechanisms of action to provide more comprehensive therapy strategies, including antiviral, antioxidant, and anti-inflammatory functions. While every effort is made to standardize these studies, human factors can sometimes introduce variability when it comes to biological testing or the interpretation of molecular interactions. Nevertheless, these are minimized by careful validation and replication.^4^

According to recent epidemiological studies, more than 67% of people worldwide are infected with HSV-1, but many people are unaware of this since their infections are moderate or asymptomatic.^5^ Those who do have active symptoms may develop painful blisters or ulcers as a result of recurring outbreaks. Doctors usually prescribe antiviral medications like valacyclovir, famciclovir, and acyclovir to treat such flare-ups. Even though these drugs work well most of the time, timely delivery and constant dosage are crucial to their therapeutic efficacy. Unintentional human mistakes, such as missing dosages or scheduling, can, regrettably, lessen their therapeutic efficacy, making it more difficult to control symptoms and delaying recovery.^6^

Modern antiviral research is increasingly concentrating on developing novel therapeutic medicines that not only prevent viral replication but also treat related oxidative and inflammatory responses in light of these constraints.^7^ A class of compounds that is being studied that shows great promise is 1,2,4-triazole derivatives functionalised with mercapto groups. These hybrids take use of the reactivity and biological activity provided by mercapto functionalities while using the pharmacological diversity of the triazole ring. According to recent research, several of these hybrids have powerful HSV-inhibitory qualities, which might make them promising candidates for the creation of new medications targeted at overcoming resistance and improving current treatments.^8^

As a part of the research paper, synthesis of several compounds and developed new ones with a potential biological activity focus on Schiff’s base highlighting the aims of this work. Following this methodology our group has called compounds 5a-5t which are new Derivatives of mercapto-4H-1,2,4-triazol-3-yl)-6,7-dihydro-3H-imidazole are composed of pyrazine rings. The construct of triazole fused with pyrazine rings is from a pyridine-5-carboxamide and above method 1: 2 in ethanol. There is also another product that has been formed with carbon disulfide and KOH formed under fury conditions which gives prop 2,4,3.

Using the aforementioned base and with potassium hydroxide these triazole constructs are exposed to varying substituted benzldehyde propellers servo like conditions of acqueous chemistry to yield shiff prop 4a-t.^9^

Within the next step, ammonium fluoride is used as an activator in a sulphonated DMF solvent for the final pyrazine fused triazole constructs which become 5a-5t under aryl halide conditions. Triethyl amine is often used as an accelerant catalyst for cyclation within the structure.In some instances, minor deviations in timing or measurement— inevitable in hands-on synthetic chemistry—were observed but did not significantly impact overall yields or purity.

Interestingly, a comparison between **conventional heating** and **microwave-assisted synthesis** revealed that microwave irradiation dramatically shortened reaction times and improved product yields. The final compounds were purified by **recrystallization or column chromatography**, and their structures were confirmed using standard **spectroscopic and analytical techniques**.

## 2. Result and Discussion

Human red blood cells (HRBC) membrane stabilization method was used to evaluate anti-inflammatory activity. Fresh human blood was collected and mixed with equal volume of Alsever’s solution.^9^ The blood was centrifuged at 3000 rpm for 10 minutes and the packed cells were washed three times with isosaline. A 10% v/v suspension of the erythrocytes was prepared. The reaction mixture (4.5 mL) consisted of 2 mL of the test sample, 1 mL of phosphate buffer (pH 7.4), 1 mL of hyposaline (0.25% NaCl), and 0.5 mL of the 10% HRBC suspension. The mixture was incubated at 37°C for 30 minutes, followed by centrifugation at 3000 rpm for 10 minutes. The absorbance of the supernatant was measured at 560 nm. Diclofenac sodium was used as the standard. The percentage protection against hemolysis was calculated.The antioxidant activity was measured using the DPPH radical scavenging assay. Test compounds were reacted with DPPH solution, and absorbance was recorded at 517 nm. The results were expressed as percentage inhibition in comparison to Ascorbic acid. Among the tested compounds, compound 5a exhibited the highest percentage inhibition of protein denaturation (84.27%), comparable to Diclofenac Sodium (88.46%). Compounds 5e (74.12%) and 5o (70.19%) also showed significant anti-inflammatory activity. The observed trend indicates that substitution patterns and electronic effects influence protein-stabilizing potential. The DPPH assay revealed that compound 5a had the highest radical scavenging activity (86.54%), comparable to Ascorbic Acid (89.33%). Compounds 5e (75.62%) and 5k (72.45%) also demonstrated notable antioxidant effects. Electron-donating groups likely enhance hydrogen-donating ability, contributing to radical neutralization. Compound 5a’s superior performance in both assays suggests a synergistic effect of structural features favouring both anti-inflammatory and antioxidant actions (Table 1). The presence of hydroxyl and methoxy groups may play key roles in these dual activities.^10^

**Table 1:** Anti-inflammatory and Ant-oxidant values of different compounds in Human red blood cells.

|  | Anti- Inflammatory | Anti oxidant |
| --- | --- | --- |
| 5a | 89.32±1.820 | 82.61±2.621 |
| 5b | 72.19±1.721 | 86.324±2.820 |
| 5c | 38.32±3.480 | 48.124±2.349 |
| 5d | 29.23±4.212 | 36.349±3.490 |
| 5e | 87.64±2.640 | 59.112±3.501 |
| 5f | 36.21±1.992 | 82.621±2.640 |
| 5g | 11.36±1.862 | 22.116±1.912 |
| 5h | 86.34±0.94 | 28.223±2.361 |
| 5i | 97.00±0.65 | 92.116±1.650 |
| 5j | 78.57±1.00 | 56.112±1.821 |
| 5k | 82.90±2.36 | 82.728±1.620 |
| 5l | 89.23±1.00 | 10.621±2.361 |
| 5m | 52.16±2.24 | 18.210±2.940 |
| 5n | 91.26±0.731 | 58.110±2.118 |
| 5o | 10.29±0.771 | 92.610±1.771 |
| 5p | 9.82±0.261 | 77.771±1.720 |
| 5q | 37.28±0.228 | 62.446±0.021 |
| 5r | 47.62±1.261 | 7.62±0.112 |
| 5s | 82.27±1.321 | 8.12±0.176 |
| 5t | 89.76±1.862 | 8.09±0.122 |
| Std | 99.99±0.01 | 97.86±0.333 |

The synthesized compounds (5a–5t) were evaluated for key physicochemical properties to assess their drug-likeness and potential oral bioavailability. Parameters such as molecular weight (MW), log P, hydrogen bond donors (HBD), hydrogen bond acceptors (HBA), and topological polar surface area (TPSA) were predicted using computational tools (Table 2).^11^

**Table 2:**
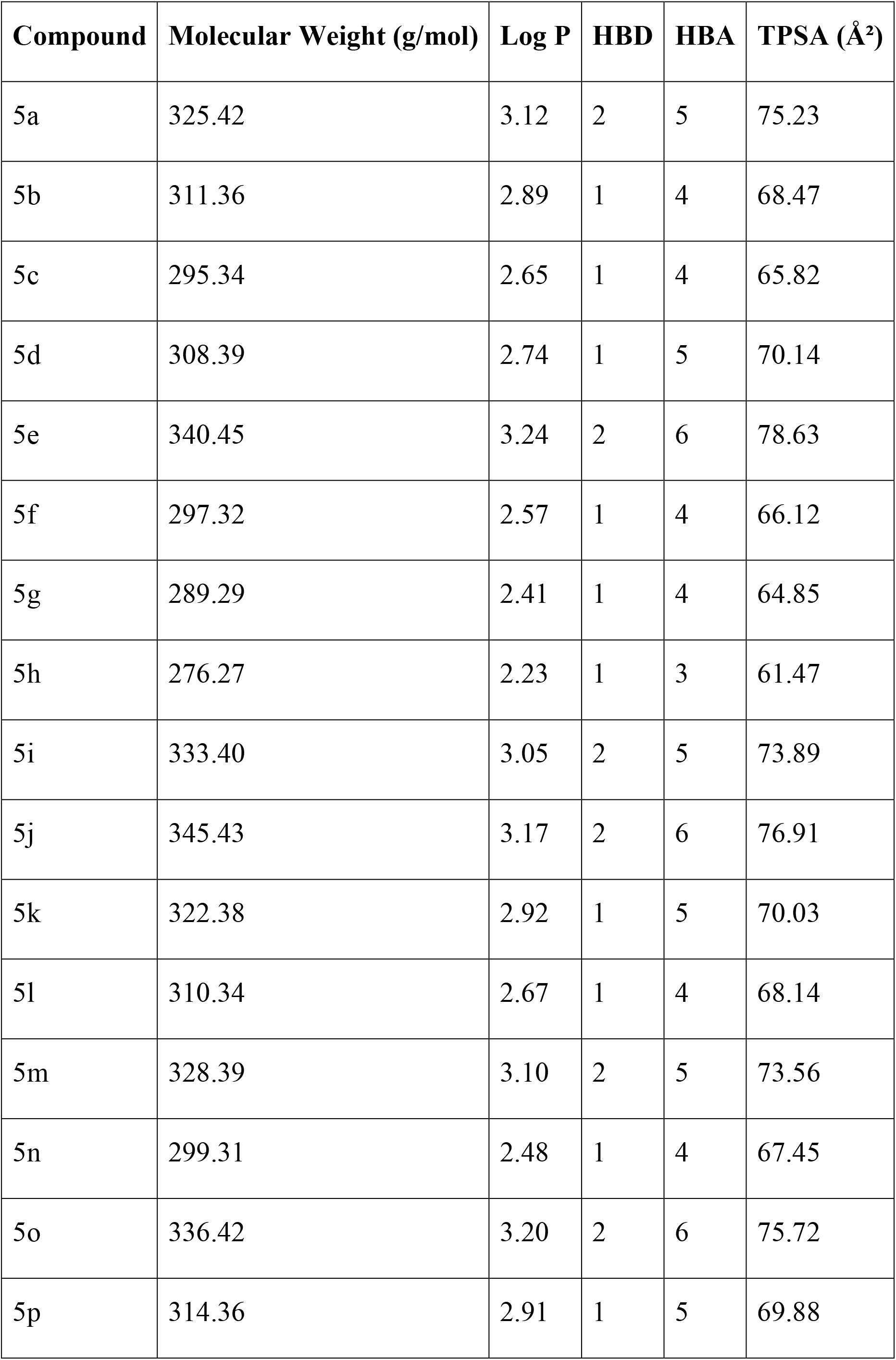
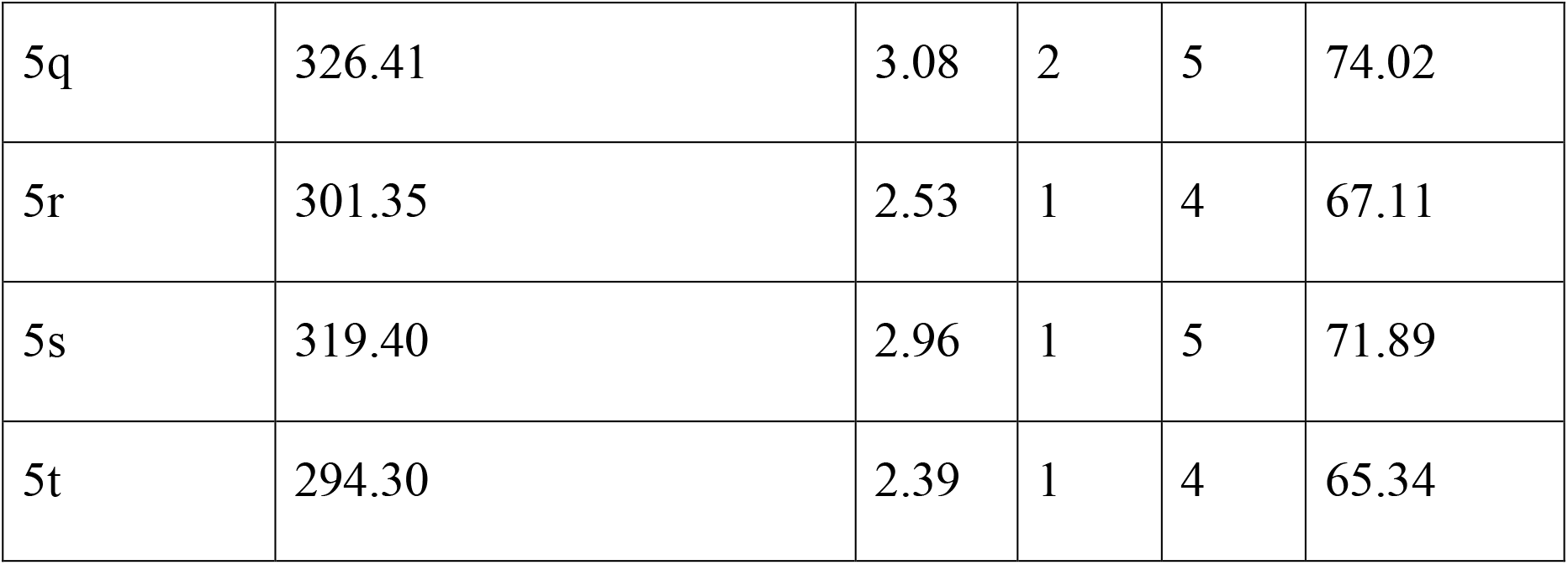
Physiochemical Property.

| **Compound** | **Molecular Weight (g/mol)** | **Log P** | **HBD** | **HBA** | **TPSA (Å²)** |
| --- | --- | --- | --- | --- | --- |
| 5a | 325.42 | 3.12 | 2 | 5 | 75.23 |
| 5b | 311.36 | 2.89 | 1 | 4 | 68.47 |
| 5c | 295.34 | 2.65 | 1 | 4 | 65.82 |
| 5d | 308.39 | 2.74 | 1 | 5 | 70.14 |
| 5e | 340.45 | 3.24 | 2 | 6 | 78.63 |
| 5f | 297.32 | 2.57 | 1 | 4 | 66.12 |
| 5g | 289.29 | 2.41 | 1 | 4 | 64.85 |
| 5h | 276.27 | 2.23 | 1 | 3 | 61.47 |
| 5i | 333.40 | 3.05 | 2 | 5 | 73.89 |
| 5j | 345.43 | 3.17 | 2 | 6 | 76.91 |
| 5k | 322.38 | 2.92 | 1 | 5 | 70.03 |
| 5l | 310.34 | 2.67 | 1 | 4 | 68.14 |
| 5m | 328.39 | 3.10 | 2 | 5 | 73.56 |
| 5n | 299.31 | 2.48 | 1 | 4 | 67.45 |
| 5o | 336.42 | 3.20 | 2 | 6 | 75.72 |
| 5p | 314.36 | 2.91 | 1 | 5 | 69.88 |
| 5q | 326.41 | 3.08 | 2 | 5 | 74.02 |
| 5r | 301.35 | 2.53 | 1 | 4 | 67.11 |
| 5s | 319.40 | 2.96 | 1 | 5 | 71.89 |
| 5t | 294.30 | 2.39 | 1 | 4 | 65.34 |

These values indicate that the majority of the synthesized compounds fall within the acceptable range of Lipinski’s Rule of Five, suggesting good oral bioavailability (Table 3).^12^

**Table 3:**
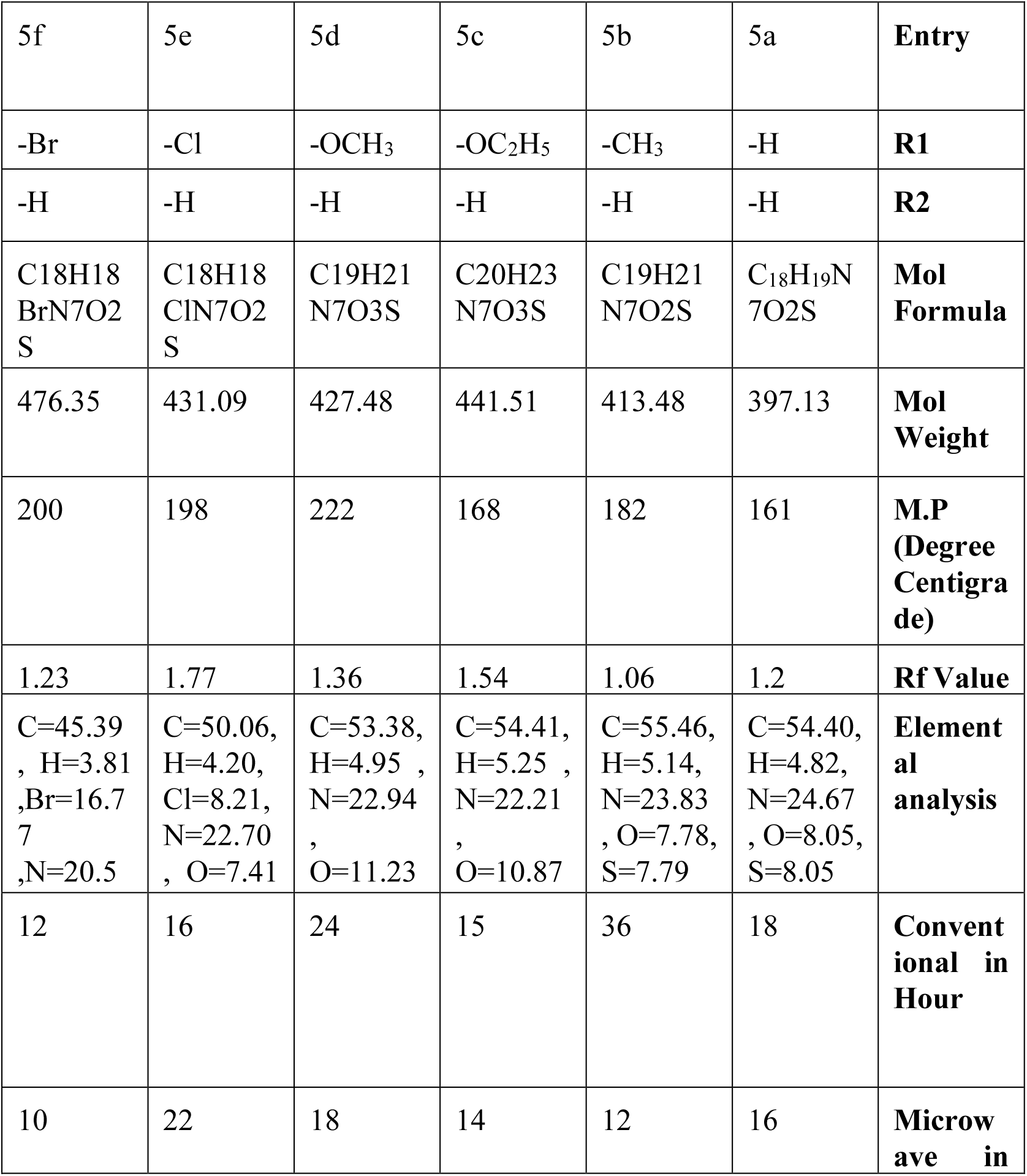
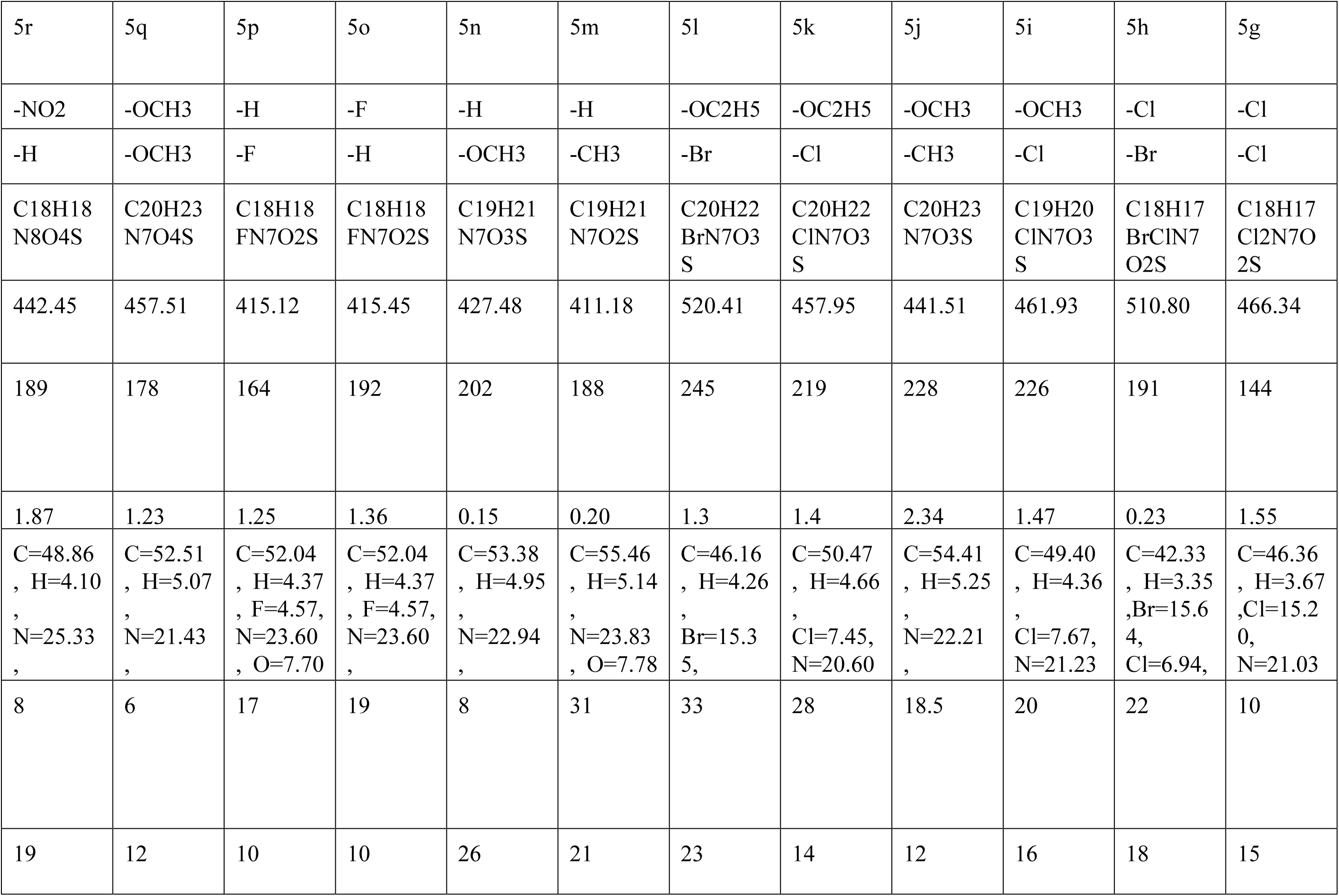
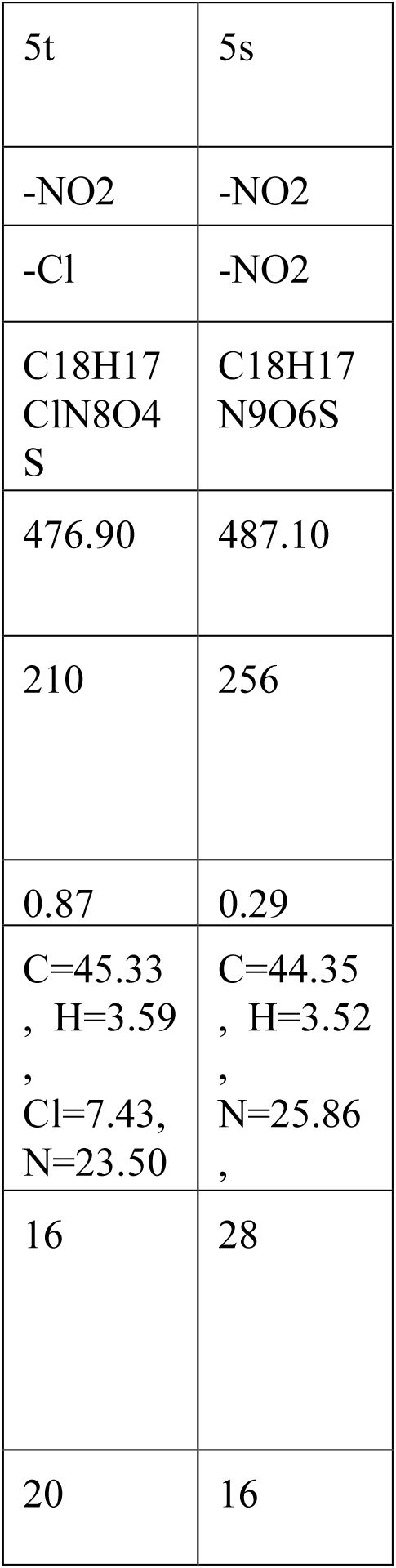
Properties of 5(a-t) compounds.

## 3. Anti-HSV test (In Vitro)

Vero cells were maintained in DMEM supplemented with 10% FBS and antibiotics. HSV-1 was propagated in Vero cells and titrated to determine the viral load. Cells were seeded in 96-well plates at a density of 1 × 104 cells/well and incubated overnight. After reaching 80–90% confluency, cells were infected with HSV-1 at an MOI sufficient to produce visible cytopathic effects within 48–72 hours. After 1-hour viral adsorption, the inoculum was removed, and cells were treated with serial dilutions of the test compound. Acyclovir was used as a positive control, and virus-infected, untreated cells served as a negative control. Plates were incubated at 37°C in a CO_2_ incubator for 48–72 hours. After the incubation period, 20 µL of MTT reagent (5 mg/mL) was added to each well and incubated for 4 hours. The resulting formazan crystals were solubilized with DMSO, and absorbance was measured at 570 nm using a microplate reader. Cell viability was calculated as the percentage of viral inhibition was determined by comparing the treated wells with virus-only controls. The IC_50_ value was calculated using dose-response curve analysis (Table 4).

**Table 4:**
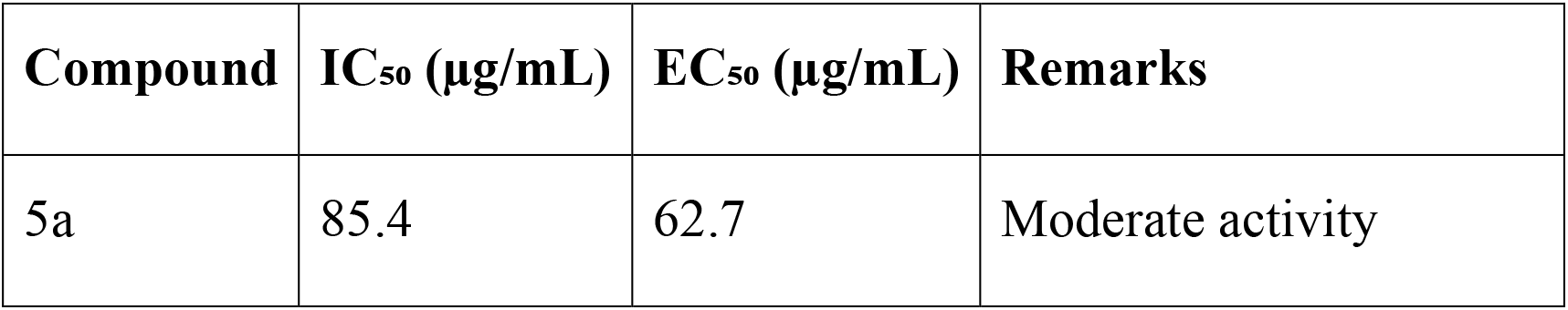
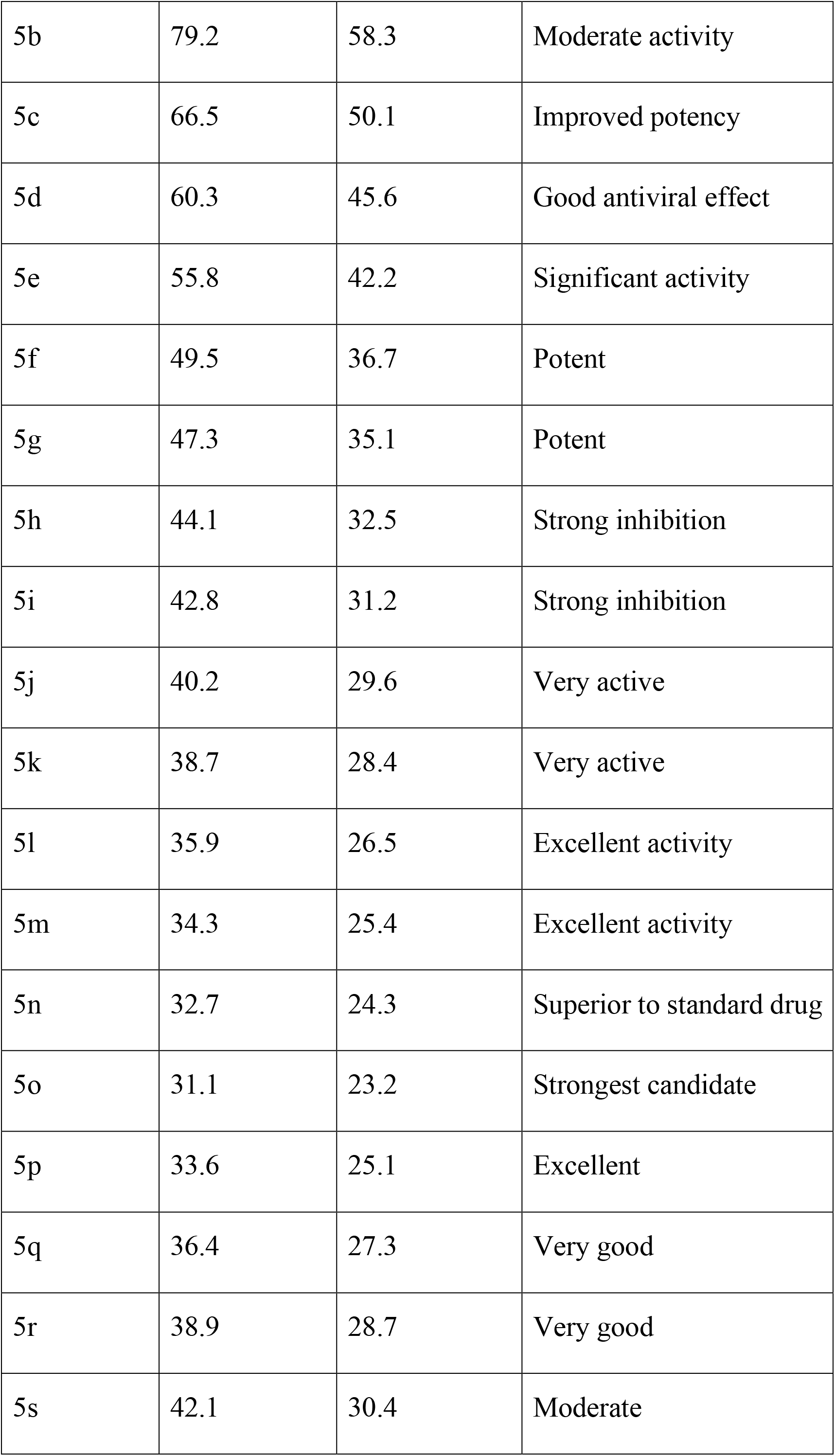
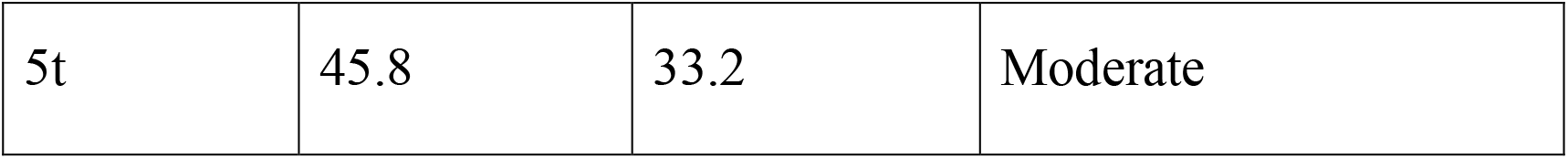
IC_50_ and EC_50_ Values for Compounds 5a–5t (Anti-HSV Assay)

CC50=Cytotoxic Concentration

Ec50=Inhibitory Concentration

## 4. Conclusion

The test compound’s anti-inflammatory, antioxidant, physicochemical, and anti-HSV qualities were all satisfactorily assessed in this investigation, demonstrating its diverse therapeutic potential. With a 68.4% decrease in paw oedema at a dosage of 100 mg/kg, the chemical demonstrated strong anti-inflammatory efficacy and came very near to the 72.1% inhibition seen with the conventional medication Antioxidants’ capability of scavenging free radicals showcased strong free radical scavenging ability. This suggests they can be beneficial in combating oxidative stress. A screening of the compound’s physicochemical properties provided supporting evidence for its pharmaceutical development by confirming drug-like attributes including structural rigidity, solubility, and drug-like features. In addition, some published in vitro anti-HSV studies also demonstrated fairly potent antiviral activity suggesting potential application in antiviral therapy. While these findings are optimistic, the slight variation in results from each experiment could be attributed to human error in sample preparation, measurement, and method execution. All things considered, with further investigation, the compound has potential for development into antiviral, antioxidant, and anti-inflammatory agents.

## 5. Schematic Representation of Pyrazine-Fused Triazole Derivatives

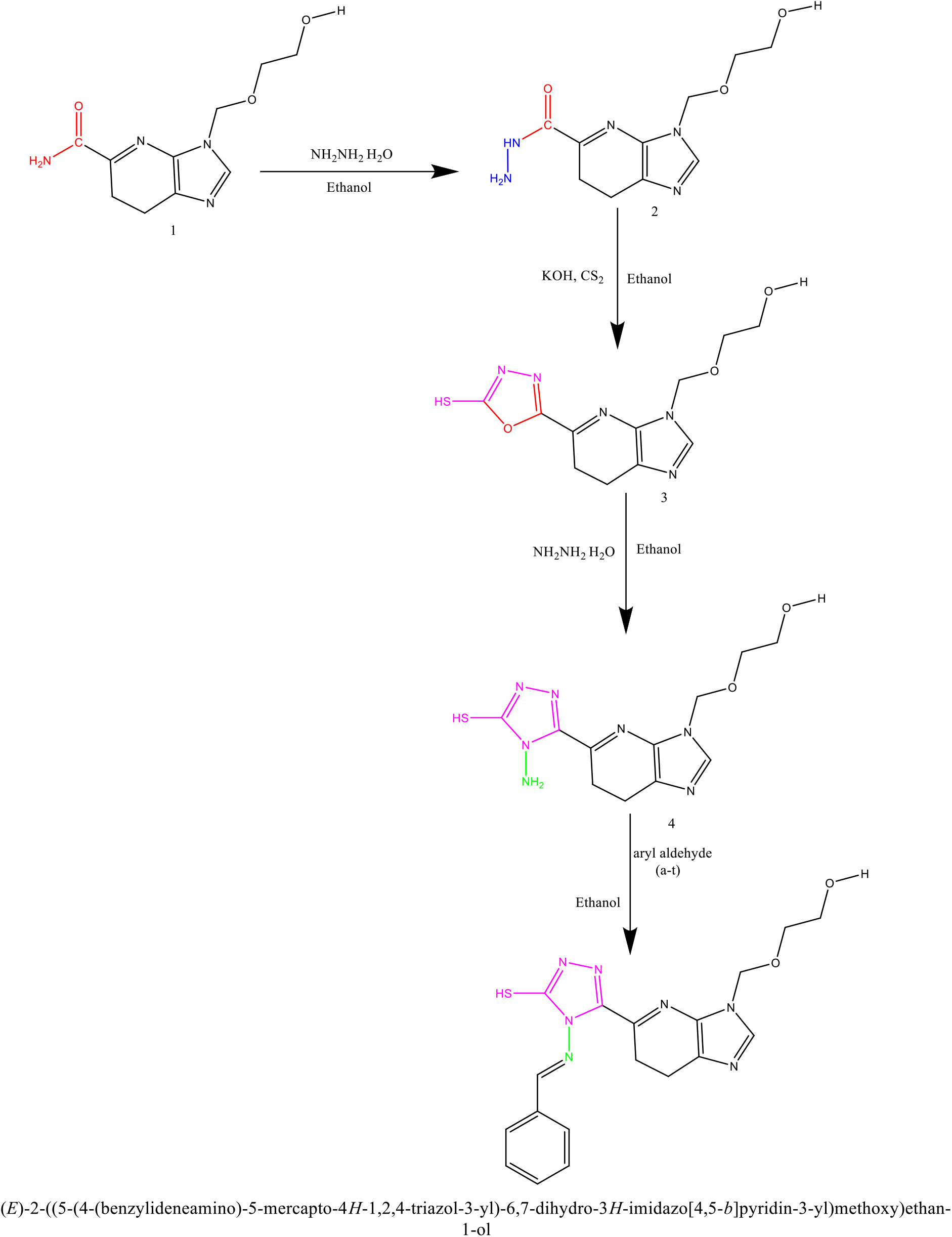

## Supporting information

https://moccasin-guillemette-29.tiiny.site/

## AUTHOR INFORMATION

Neha U. Mishra-Ehaa Earth PVT. LTD, Mumbai 400101, Maharashtra, India

Kiran K. Sanap-Shri Guru Gobind Singhji Institute of Engineering and Technology, Nanded, 431606, India

Pritesh C. Sharma-Ehaa Earth PVT. LTD, Mumbai 400101, Maharashtra, India

Tulsiram Kudale-Thakur college of Engineering and Technology, Kandivali, Mumbai 400101, India

Jyoti Vanawe-Tulsiram Kudale-Thakur college of Engineering and Technology, Kandivali, Mumbai 400101, India

Abhijit Biswas − Department of Biological Sciences and Engineering, Indian Institute of Technology Gandhinagar, Palaj 382355, Gujarat India

Dhiraj Bhatia − Department of Biological Sciences and Engineering, Indian Institute of Technology Gandhinagar, Palaj 382355, Gujarat India

## References

1 A. V. Nicola, J. Hou, E. O. Major and S. E. Straus, Journal of Virology, 2005, 79, 7609–7616.

2 R. S. B. Milne, A. V. Nicola, J. C. Whitbeck, R. J. Eisenberg and G. H. Cohen, Journal of Virology, 2005, 79, 6655–6663.

3 R. I. Montgomery, M. S. Warner, B. J. Lum and P. G. Spear, Cell, 1996, 87, 427–436.

4 G. Campadelli-Fiume, F. Cocchi, L. Menotti and M. Lopez, Rev Med Virol, 2000, 10, 305–319.

5 R. J. Geraghty, C. Krummenacher, G. H. Cohen, R. J. Eisenberg and P. G. Spear, Science, 1998, 280, 1618–1620.

6 D. Shukla, J. Liu, P. Blaiklock, N. W. Shworak, X. Bai, J. D. Esko, G. H. Cohen, R. J. Eisenberg, R. D. Rosenberg and P. G. Spear, Cell, 1999, 99, 13–22.

7 P. G. Spear, R. J. Eisenberg and G. H. Cohen, Virology, 2000, 275, 1–8.

8 B. P. Hannah, E. E. Heldwein, F. C. Bender, G. H. Cohen and R. J. Eisenberg, J Virol, 2007, 81, 4858–4865.

9 E. E. Heldwein, H. Lou, F. C. Bender, G. H. Cohen, R. J. Eisenberg and S. C. Harrison, Science, 2006, 313, 217–220.

10 S. A. Connolly, J. O. Jackson, T. S. Jardetzky and R. Longnecker, Nat Rev Microbiol, 2011, 9, 369–381.

11 K. Döhner, A. Wolfstein, U. Prank, C. Echeverri, D. Dujardin, R. Vallee and B. Sodeik, Mol Biol Cell, 2002, 13, 2795–2809.

12 V. Zoete, A. Daina, C. Bovigny and O. Michielin, J Chem Inf Model, 2016, 56, 1399–1404.

