## Supplementary material for "Synthesis and In Vitro Assessment of Triazole-Based Compounds as Potential Inhibitors of Herpes Simplex Virus Type 1": https://moccasin-guillemette-29.tiiny.site/

### Supporting Information's:

#### 1. Step-wise Synthesis of Pyrazine-Fused Triazole Schiff Base Derivatives (5a–5t)

**Step 1: Synthesis of Triazole Intermediate**

- **Reactants:** Pyridine-5-carboxamide and hydrazine hydrate
- **Molar Ratio:** 1:2
- **Solvent:** Ethanol
- **Conditions:** Reflux for 7 hours under stirring
- The hydrazide group provides a precursor for cyclization with carbon disulfide.

**Step 2: Cyclization with Carbon Disulfide**

- **Reactants:** Hydrazide intermediate, Carbon disulfide ($CS_2$), and Potassium hydroxide (KOH)
- **Conditions:** Stirring under heat 85°C
- **Solvent/Base Medium:** alcoholic KOH
- **Reaction Type:** Cyclization and sulfur incorporation
- Introduces the biologically active triazole core with a thiol functional group.

**Step 3: Schiff Base Formation (Compounds 4a–4t)**

- **Reactants:** Triazole-thiol intermediate (from Step 2) and substituted benzaldehydes
- **Conditions:** Aqueous basic medium, reflux 55°C
- **Catalyst/Base:** Potassium hydroxide (KOH)
- **Reaction Type:** Nucleophilic addition followed by condensation (C=N bond formation)

**Step 4: Final Pyrazine Fused Cyclization (Compounds 5a–5t)**

- **Reactants:** Schiff base derivatives (4a–4t), aryl halides
- **Solvent:** Sulphonated Dimethylformamide (DMF)
- **Additives:**
  - **Ammonium fluoride ($NH_4F$):** Used as an activator
  - **Triethylamine ($Et_3N$):** Used as a catalyst and base to promote cyclization
- **Conditions:** Heating under controlled temperature (145°C)
- **Reaction Type:** Cyclization and arylation under nucleophilic substitution conditions

Introduction of fused pyrazine ring via electrophilic substitution for enhanced rigidity and biological relevance.

**Step 4: Final Cyclization to Form Pyrazine-Fused Triazole Derivatives (5a–5t) Using Microwave-Assisted Synthesis (Anton Paar Reactor)**

- **Reactants:** Schiff base derivatives (4a–4t) and appropriate aryl halides
- **Solvent:** Sulphonated Dimethylformamide (DMF)
- **Catalyst/Base:**
    - **Ammonium fluoride ($NH_4F$):** Used as an activator to enhance nucleophilic substitution
    - **Triethylamine ($Et_3N$):** Serves as a base and cyclization catalyst
- **Equipment: Anton Paar Microwave Reactor**
    - **Mode:** Closed-vessel microwave-assisted synthesis
    - **Temperature:** 120 °C
    - **Time:** 15–30 minutes
    - Automatic internal pressure monitoring
    - **Power Input:** Optimized between 100–200 W depending on scale and reactivity

**2. Schematics of Pyrazine-Fused Triazole Derivatives (5a–5t)**

| | |
|---|---|
| 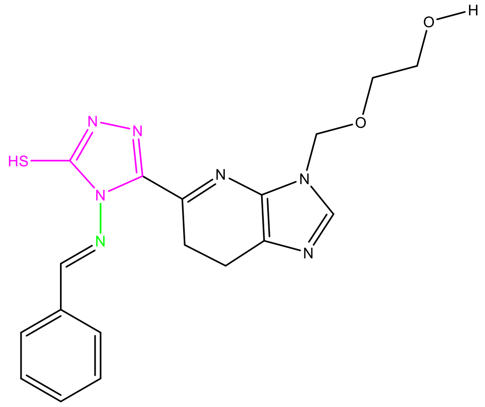 | 5a |
| 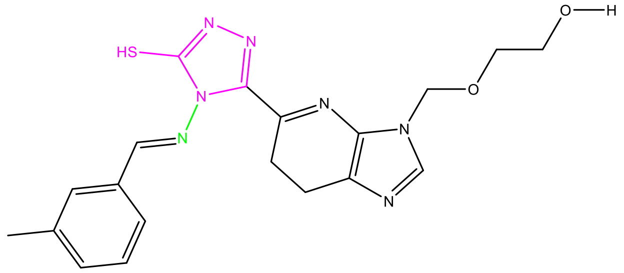 | 5b |

| | |
|---|---|
| 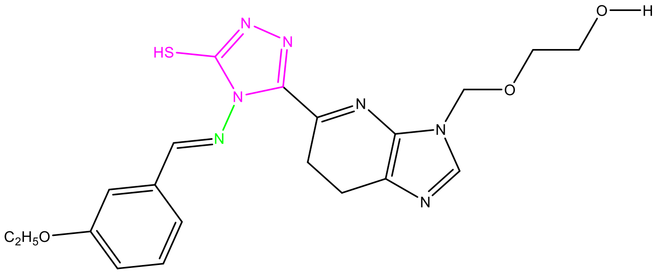 | 5c |
| 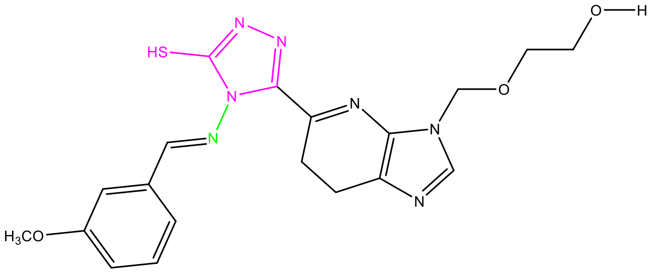 | 5d |
| 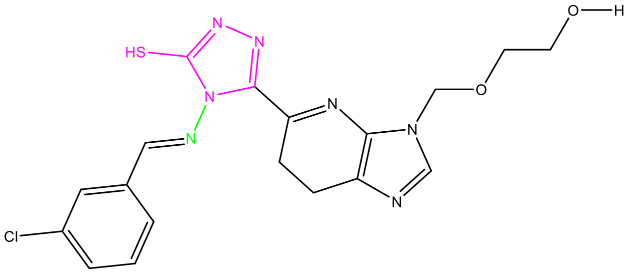 | 5e |
| 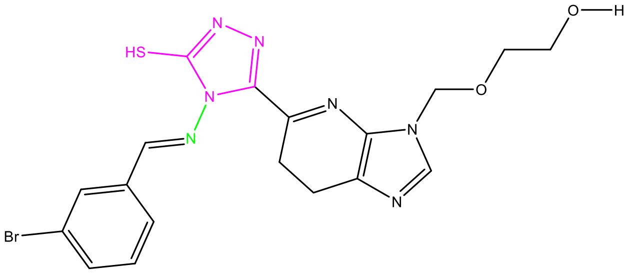 | 5f |
| 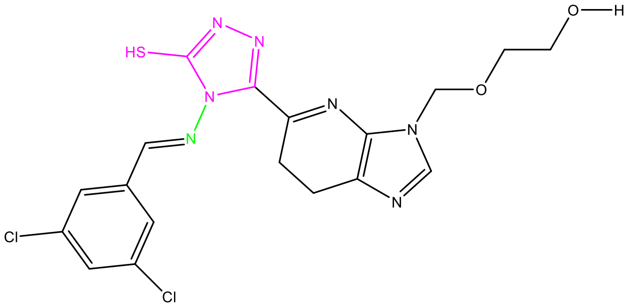 | 5g |
| 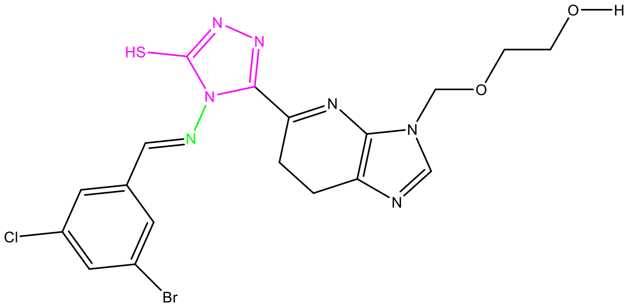 | 5h |

| | |
|---|---|
| 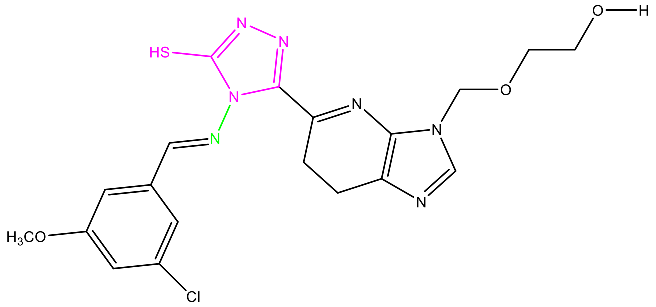 | 5i |
| 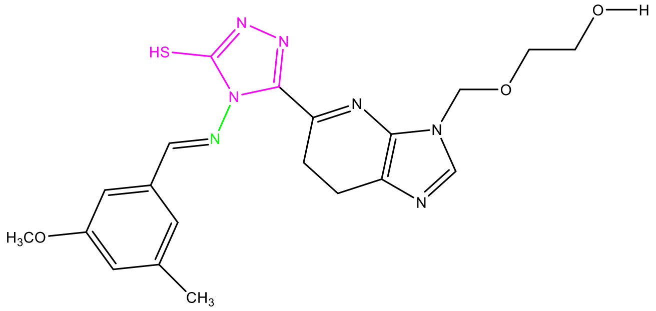 | 5j |
| 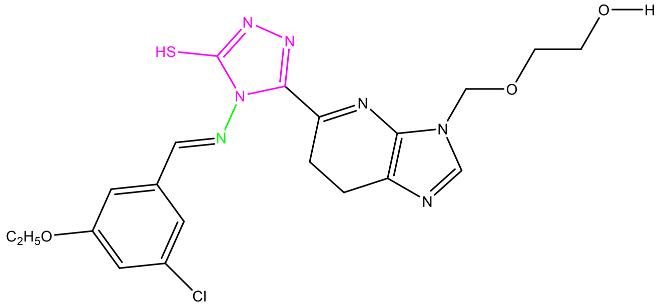 | 5k |
| 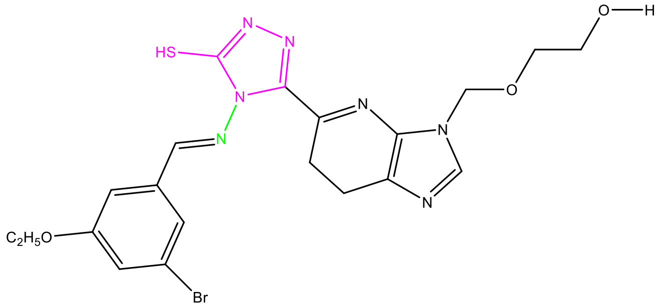 | 5l |
| 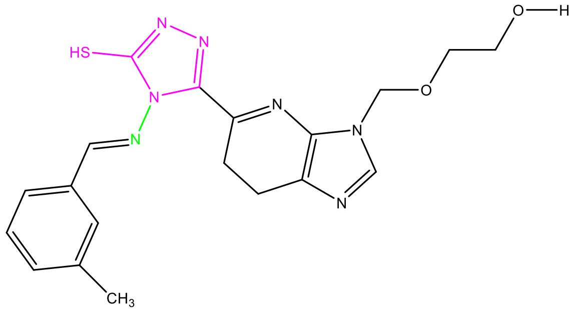 | 5m |

| | |
|---|---|
| 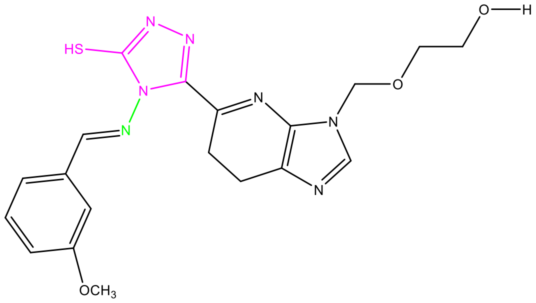 | 5n |
| 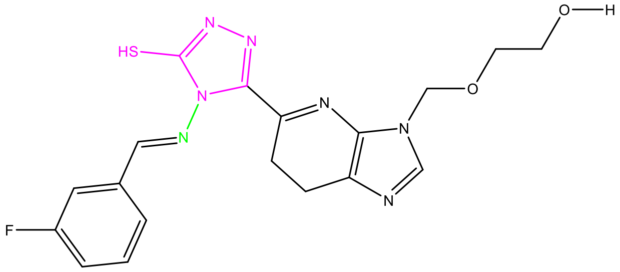 | 5o |
| 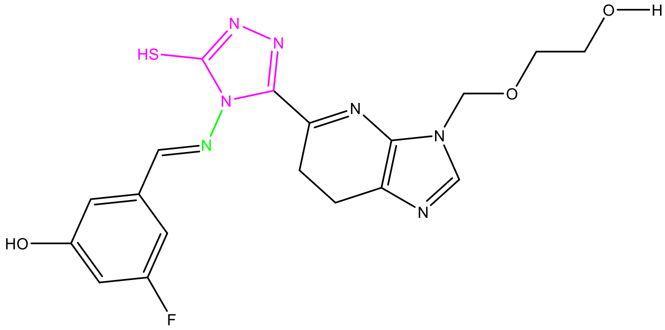 | 5p |
| 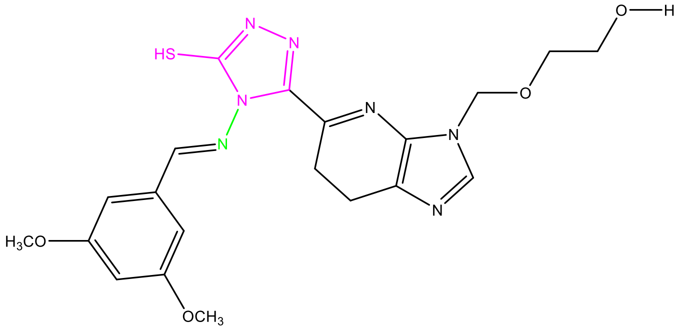 | 5q |
| 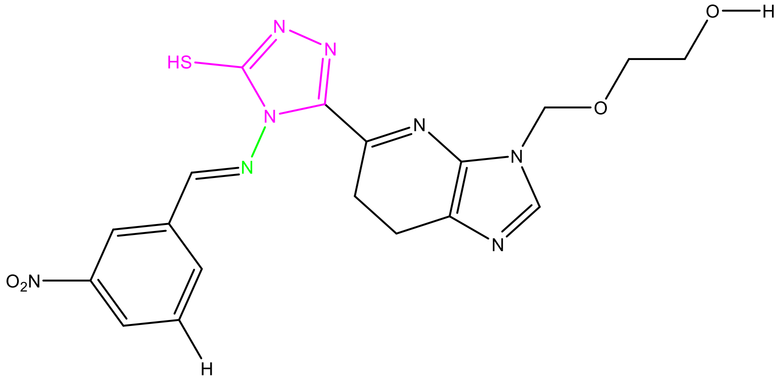 | 5r |

| | |
|---|---|
| 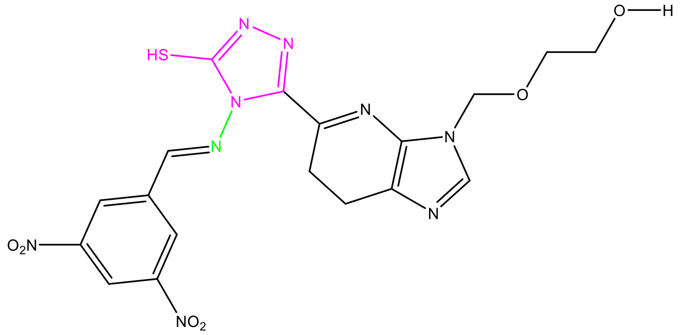 | 5s |
| 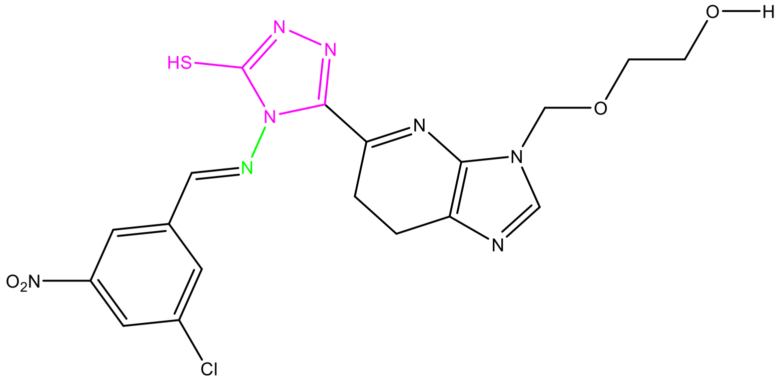 | 5t |
